# Comparison of stimulus-evoked BOLD responses in human and monkey visual cortex

**DOI:** 10.1101/345330

**Authors:** Gaurav H. Patel, Alexander L. Cohen, Justin T. Baker, Lawrence H. Snyder, Maurizio Corbetta

## Abstract

We characterized the blood oxygenation level dependent (BOLD) signal in humans and macaque monkeys by comparing the response in visual cortex to a single checkerboard or two checkerboards, spaced 1.5, 3.0, or 4.5 s apart. We found that the magnitude and shape of the BOLD response to a single checkerboard was similar in the two species. In addition, we found that the BOLD responses summed similarly, and that at an inter-stimulus interval (ISI) of 4.5 sec BOLD summation was nearly linear in both species. When comparing the ratio of the amplitude of the response to the second checkerboard at the 4.5 sec ISI with that of the single checkerboard between subjects in both species, the results from both monkey subjects fell within one standard deviation of the mean human results (human mean (n=12): .95 +/− .31 second/single response amplitude; monkey 1: 1.16; monkey 2: .86). At the shorter ISIs, both species demonstrated increased suppression of the BOLD response to the second checkerboard. These findings indicate that the magnitude of the BOLD response to events separated by 4.5 seconds can be accurately measured in and compared between human and monkey visual cortex.

## Introduction

Recent developments in functional magnetic resonance imaging (fMRI) have made feasible the *in vivo* visualization of neural activity in awake behaving macaques (Vanduffel et al., 2001; Fize et al., 2003; Tsao et al., 2003a; Tsao et al., 2003b; Denys et al., 2004a; Denys et al., 2004b; Koyama et al., 2004; Nelissen et al., 2005; Pinsk et al., 2005; Sawamura et al., 2005; Baker et al., 2006). This advancement is significant for several reasons. First, monkey fMRI provides a relatively easy way to obtain a whole-brain view of task-evoked neural activity. This map of activity can be used to guide more in-depth explorations of neural activity using electrode-based methods, such as single or multi-unit recordings, which may in turn help us better understand models of cognition developed with human fMRI data. Second, fMRI allows for the direct comparison of neural activity in macaques and humans performing the same task under similar conditions, which may provide a better understanding of the evolutionary relationships between brain function and organization in the two species (Logothetis et al., 2001; Orban, 2002; Logothetis, 2003; Orban et al., 2004; Sereno and Tootell, 2005).

Functional MRI measures local changes in the blood oxygen level dependent (BOLD) signal, which are correlated with changes in local neural activity (Logothetis et al 2001). A critical issue for performing parallel fMRI studies in human and non-human primates is whether the spatial and temporal characteristics of the BOLD response are similar in the two species. In humans, the onset of the BOLD response lags behind the onset of a stimulus by about 2 seconds (sec), and the response does not return to baseline until about 16-18 sec after stimulus offset, even though the neural response may last only a few hundred milliseconds (Boynton et al., 1996; Buxton, 2002; Buxton et al., 2004). The human BOLD response to a transient stimulus can be accurately modeled using a gamma function (Friston et al., 1995; Boynton et al., 1996; Aguirre et al., 1998).

Recent studies have characterized some aspects of hemodynamic response in macaques. Logothetis and colleagues (Logothetis et al., 1999; Logothetis et al., 2001; Logothetis et al., 2002) have shown that the spatiotemporal features of the BOLD signal in macaque visual cortex under anesthesia was similar to that of humans. Vanduffel *et al.* have demonstrated similar time-courses in awake behaving macaques (Vanduffel et al., 2001). Leite and coauthors modeled the BOLD response to 2-4s of visual stimulation in awake behaving macaques using exponential functions, and demonstrated qualitative similarities between human and macaque BOLD responses (Leite et al., 2002; Leite and Mandeville, 2006). Because of these results, all of the fMRI studies comparing humans and macaques to-date have assumed that the underlying physiology of the stimulus-evoked BOLD responses in awake behaving macaques was similar enough to that of humans to use the same analysis methods and assumptions (Vanduffel et al., 2001; Koyama et al., 2004; Pinsk et al., 2005; Baker et al., 2006).

The validity of this assumption may not be critical when the goal is simply to identify the presence of significant task-evoked activity in each species. However, the accuracy of this assumption is important for inter-species comparisons of response modulation to different task variables and across brain regions. While the above studies suggest a qualitative similarity between human and macaque BOLD responses, no quantitative comparison between the two species has been performed. If the parameters of the hemodynamic response differ between the two species, then the use of the same assumed hemodynamic response function to generate statistical maps in each species could lead to erroneous results.

Another critical issue is whether and to what extent BOLD responses in humans and monkeys add linearly in time. Rapid event-related designs allow for the separation of specific sensory, cognitive, and motor events by separating each of these event types in time, while maximizing signal to noise by minimizing this inter-event spacing as much as possible. However, due to the sluggishness of the neurovascular transformation, the BOLD response to one event overlaps with that of the following events, making it impossible to resolve the single-event response just by examining the BOLD signal time-course (Bandettini et al., 1992; Ogawa et al., 1992). However, if the overlapping BOLD responses add linearly, then a general linear model (GLM) can be used to disentangle the responses to individual events (Friston et al., 1995; Dale and Buckner, 1997).

In human subjects, Dale and Buckner showed that BOLD responses to transient visual stimuli added linearly as long as the stimulus events were separated by inter-stimulus intervals (ISIs) of at least 5 seconds (Dale and Buckner, 1997). In other words, the measured response to both events can be modeled as the sum of the BOLD response to each stimulus. However, linearity breaks down at ISIs less than 5 seconds. This breakdown appears as a decreased estimated response to the second stimulus. The amplitude of the second response decreases with decreasing ISIs (Dale and Buckner, 1997; Huettel and McCarthy, 2000; Miezin et al., 2000; Boynton and Finney, 2003). Non-linearity in BOLD response summation may produce different estimation errors in different experimental conditions. In designs in which there is a variable interval between events, departures from linearity can lead to reduced power in detecting BOLD signals due to the increased variation in the response magnitude across trials. In designs in which proper counterbalancing between different event-types is not possible, such as working memory experiments in which the cue and target responses are modeled separately, non-linearities can lead to incorrect estimation of the response magnitudes (Aguirre et al., 1998).

There are many possible sources of BOLD response non-linearities—they may be purely vascular in nature, purely neural, or in differences in the neurovascular coupling that underlies the BOLD effect—but because so little is known about their origin, it is entirely possible that they might differ across species (Friston et al., 2000; Buxton et al., 2004; Logothetis and Wandell, 2004). To date this important issue has not been directly addressed in the growing field of monkey fMRI. The goals of the present study were (1) to compare the parameters of the BOLD response to a single visual event between macaques and humans and (2) to compare the relationship between BOLD response linearity and ISI length in humans and monkeys.

## Materials and Methods

### Animal Subjects, Surgery, and Experimental Setup

Two male monkeys (Macaca fascicularis, 3-5 kg) were used in accordance with Washington University and NIH guidelines. Prior to training, surgery was performed in aseptic conditions under isofluorane anesthesia to implant a head restraint device. The head restraint was constructed from polyetheretherketone (PEEK) and was anchored to the skull with dental acrylic and 10-14 ceramic screws (4 mm diameter, Thomas Recording GmbH, Germany).

The monkeys were trained to fixate in a setup that simulated the fMRI scanner environment. In both the training setup and scanner, the animal sat horizontally in a “sphinx” position inside of a cylindrical monkey chair with its head rigidly fixed by the restraint device to a head holder on the chair (Primatrix Inc., Melrose, MA). An LCD projector (Sharp USA, Montclair, NJ) was used to present visual stimuli on a screen that was positioned at the end of the bore, 75 cm from the monkeys’ eyes. A flexible plastic waterspout was positioned near the animal’s mouth for delivery of liquid rewards. Eye movements were monitored by an infrared tracking system, using an IR illuminator positioned 5-6 cm from the monkey’s left eye (ISCAN Inc., Melrose, MA). The eye position coordinates were relayed to the behavioral control system, which consisted of two linked computers running custom software. In addition to recording eye position, this system recorded scanner synchronization signals, presented visual stimuli, and delivered rewards. For further details about the experimental setup, see Baker *et al.* (Baker et al., 2006).

### Human Subjects and Experimental Setup

Thirteen human subjects (ages 22-26, 5 female) were recruited from the university community. The subjects had normal or corrected-to-normal vision. Each subject was provided informed consent according to the approved guidelines of Washington University and the NIH, and each was paid for their participation. One subject’s data was later excluded from the analysis due to excessive movement during BOLD data acquisition.

The conditions of the human scanning sessions mimicked those of the monkey scanning sessions, though several differences were necessary. Subjects lay supine in the scanner, and a mirror positioned above their eyes reflected the screen at the end of the bore, 81 cm away. The fiber-optic IR illuminator was mounted on one of the struts of the MR head-coil, and illuminated the subjects’ eye via the mirror above the subjects’ eyes (5-6 cm distance between eye and illuminator). An IR sensitive camera was mounted at the end of the bore, and monitored the subjects’ eyes via the same mirror. The same behavioral control program used for the monkeys was also used for the humans; no liquid reward was given to the humans.

### Display and Task

The same display and task were used for both the human and monkey experiments. At the beginning of a trial, a white circle (0.5¼ diameter) appeared in the center of the screen, and remained present throughout the duration of a scan. The subject’s task was to visually fixate the white circle. The visual stimulus was a circular black-and-white flickering checkerboard reversing contrast at 8Hz. Because the distance between the screen and the subject’s eye differed by 6cm between the monkeys and humans, the visual angle subtended by the checkerboard was slightly different for the two species: the checkerboard covered the visual field from 1.8 ° to 12.9° eccentricity for the monkeys and 1.7° to 12.0° for the humans. The flickering checkerboard was displayed for 0.5 sec (4 contrast reversals). Four types of trials were randomly interleaved: a single presentation of the checkerboard, or two checkerboards separated by an inter-stimulus interval of 1.5, 3.0, or 4.5 sec (**Figure 1A, B**). Each single or double stimulus presentation was followed by a 17-20 sec period of fixation during the inter-trial interval. The monkeys received a small liquid reward every 2 sec (monkey 1) or 1.5 sec (monkey 2) for maintaining fixation within a .5° radius circle around the central point. These reward schedules were chosen to reduce motion artifacts from jaw movements by minimizing the number of rewards given to the monkey, while giving rewards frequently enough that they were effectively included as part of the baseline BOLD signal rather than being explicitly modeled as separate events (see **Analysis** below). The trials were not aborted for either species if the subject failed to maintain fixation.

**Figure 1:**
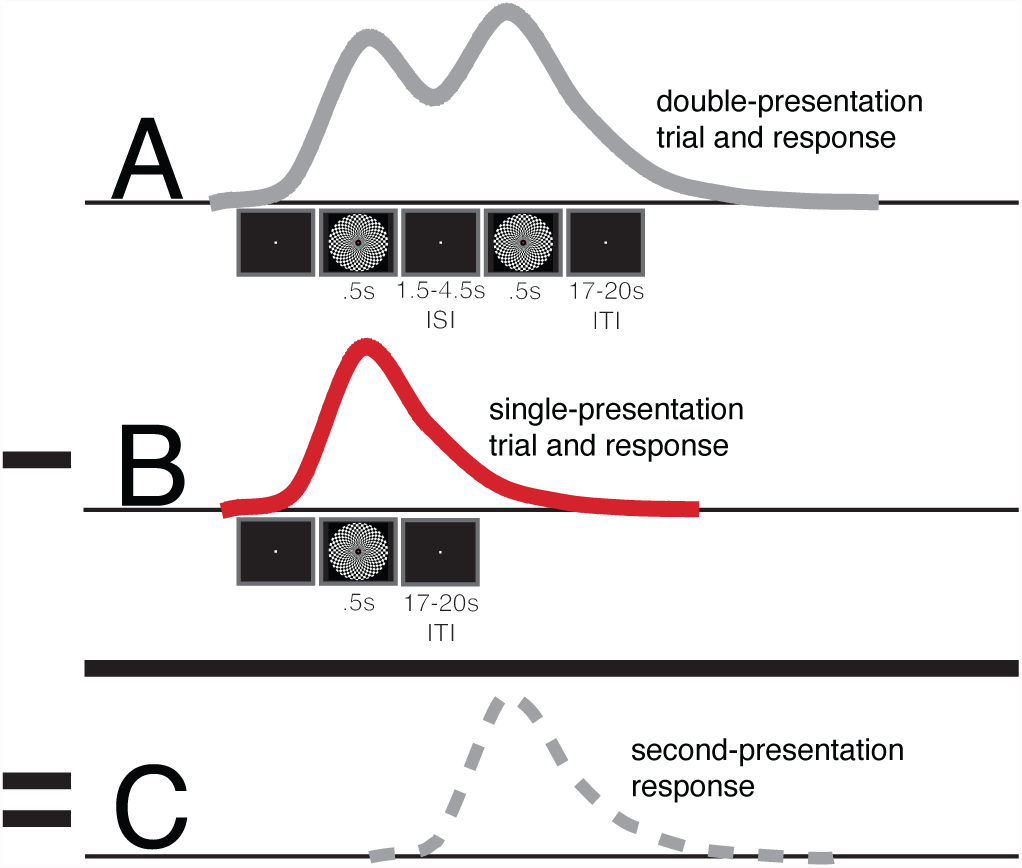
Experimental Paradigm, Hypothetical Responses, and Analysis Schematic. Subjects were run on two types of trials. In trial type **A** two checkerboards were displayed sequentially for 500ms each, separated by a 1.5, 3.0, or 4.5 sec inter-stimulus interval (ISI), and then followed by 17-20 secs of an inter-trial interval (ITI). In trial type **B** only a single checkerboard was displayed for 500ms, followed by a 17-20 sec ITI. The analysis consisted of comparing the response to the single checkerboard (**B**) with the responses to the second checkerboard at the 3 ISIs (**C**) after the first response time-course was removed by regression from the double-presentation time-course (**A**). For clarity, idealized hemodynamic responses are displayed in each panel.

### Monkey fMRI Data Collection

Functional and anatomical data in the monkey were collected in a Siemens 3T Allegra MRI scanner (Siemens Medical Solutions, Erlangen, Germany). Functional data were collected using a gradient-echo echo-planar pulse sequence sensitive to BOLD contrast (T2*) (T2* evolution time = 25 ms, flip angle = 90°) and a transmit-receive surface coil (13 cm inner diameter; Primatrix). The coil fit around each animal’s head post and was saddle-shaped to provide more extensive brain coverage as compared to a planar surface coil. Fifty-two coronal slices, each with a square field of view (96 × 96 mm, 64 × 64 base resolution, dorsal-to-ventral phase-encoding) and a thickness of 1.5 mm, were obtained using contiguous, interleaved acquisition, and a volume repetition time (TR) of 3000 ms. This scanning protocol was chosen to cover the whole brain at an isotropic spatial resolution of (1.5 mm)^3^. Forty-five to ninety volumes were acquired in each fMRI run, the first four of which were excluded from the analyses to allow for the equilibration of longitudinal magnetization. 73 BOLD runs of 90 frames each were collected during 6 sessions from Monkey 1 (number of trials per condition: single checkerboard—216; 1.5 sec ISI—195; 3.0 sec ISI— 196; 4.5 sec ISI—194), and 68 BOLD runs of 45-90 frames each were collected during 12 sessions from Monkey 2 (single checkerboard—130; 1.5 sec ISI—154; 3.0 sec ISI—145; 4.5 sec ISI—94). In order to sample the BOLD response at half-TR intervals, the stimuli were timed to occur either at the beginning of each frame or half-way through the frame (Miezin et al., 2000). Since the TR was 3.0 seconds, the effective temporal sampling interval was 1.5 seconds. The 17-20 sec ITI (see **Display and Task**, above) allowed for the BOLD response to return to baseline before the following visual stimulation event (Boynton et al., 1996; Huettel and McCarthy, 2000).

High-resolution structural images were collected in separate sessions, during which the animal was chemically restrained (10-15 mg/kg ketamine, 0.6 mg/kg xylazine, .05 mg/kg atropine). T1-weighted images were acquired using a magnetization-prepared rapid acquisition gradient-echo pulse sequence [MP-RAGE; (0.75 mm)^3^ isotropic resolution, flip angle = 7°, six acquisitions] and a volumetric transmit and receive coil (16 cm i.d.; Primatrix).

### Human fMRI Data Collection

The same scanner was also used to collect fMRI data from the human subjects. Both the functional and anatomical data were collected with a volumetric transmit and receive coil in the same session. Functional data were collected using a gradient-echo EPI sequence (256mm × 256mm, 64 × 64 base resolution, TR=3000, 39 transverse slices, 4mm slice thickness) with an effective resolution of (4mm)^3^. Twelve to fourteen 120-frame BOLD runs were collected from each subject, corresponding to ~50 trials per ISI condition per subject (ratio of trial type presentation frequency — single:1.5 sec ISI:3.0 sec ISI:4.5 sec ISI — 1:.88:.97:.94). The previously described jittering procedure was used to achieve 1/2 TR temporal sampling.

T1- and T2-weighted anatomical images were collected at the beginning of the session (T1: MP-RAGE sequence, FOV=256×256 mm^2^, 128 slices, 1 × 1 × 1.25mm^3^; T2: turbo-spin echo sequence, FOV=220×220 mm^2^, 45 slices, 1.1 × 1.1 × 3mm^3^).

### Preprocessing

In the monkey 2, BOLD runs in which there was excessive movement were excluded from preprocessing and analysis, which left 63/68 runs to be included in the data set; all 73 runs from monkey 1 were included. Every BOLD run from the 12 included human subjects was preprocessed and analyzed, and no trials in either species were excluded from the analysis for non-performance of the task. In both species, each reconstructed fMRI run produced a 4-dimensional (x, y, z, time) data set that was passed through a sequence of unsupervised processing steps using in-house software, including 1) correction for asynchronous slice acquisition using cubic spline interpolation; 2) correction of odd-even slice intensity differences resulting from the interleaved acquisition of slices; 3) 6-parameter rigid body realignment to correct for movement; 4) normalization and equalization of the intensity of each frame within a run; and 5) normalization across runs, in which each four-dimensional data set was uniformly scaled to a whole brain mode value of 1000. The data from each species was also aligned to a standard volume using a 6-parameter rigid-body realignment algorithm (Snyder, 1995). For the monkeys, the standard volume for each subject was a BOLD-EPI image that was constructed from the average of the first frame of several BOLD runs collected from the monkey during a previous study (Baker et al., 2006). For the humans, the target was a (3 mm)^3^ MPRAGE atlas image produced from the co-registration of 12 different individuals aligned to the Tailarach atlas (Talairach and Tournoux, 1988; Buckner et al., 2004).

### Analysis

First, in each subject of both species, a V1 region of interest (ROI) was defined as the conjunction of three criteria: voxels within V1, voxels that were significantly activated by the presentation of a single checkerboard, and voxels that were positively activated by the presentation of a single checkerboard. The borders of V1 were taken from the species-specific atlas available from the SumsDB (F99-UA1 for the macaques at http://sumsdb.wustl.edu:8081/sums/directory.do?id=679531, and PALS for the humans at http://sumsdb.wustl.edu:8081/sums/directory.do?dir_id=679528); these borders were used to define a surface region that was then used to create a 3-D V1 region aligned to each individuals’ anatomical image. Voxels were judged to be significantly activated in each individual if there was a significant effect (p<.01) in a one-way ANOVA(main effect of time, 6 frames). Lastly, voxels were judged to be positively activated in each individual by isolating all voxels throughout the brain that on average demonstrated activity above the baseline level 6 seconds after the single checkerboard presentation. Each subject’s ROI was composed of the voxels that passed all three criteria (see **Figure 2**); the time-courses extracted from this ROI were then used in all subsequent analyses. No assumed hemodynamic response was used in this ROI definition.

The time-courses were extracted in both species using the same single-subject methods. The BOLD data were first smoothed with a Gaussian kernel (1- voxel full-width at half maximum), and were then tested for significant visualstimulation-related activity using a general linear model (GLM). This model tested for activity evoked by each of the four trial-types (between-trial GLM, 4 regressors: 1) single checkerboard trials, or double checkerboard trials separated by 2) 1.5 sec, 3) 3.0 sec, or 4) 4.5 sec). No hemodynamic response was assumed in the model; instead, the trial-evoked responses were modeled with delta function regressors, which allowed for an estimation of the time-course of the BOLD response that was free of any shape assumption (Ollinger et al., 2001b; Ollinger et al., 2001a). Regressors for the baseline and linear trend of the time-series were also included. These regressors were then used to compute a new time-series in which the baseline and linear trend signals had been removed.

**Figure 2:**
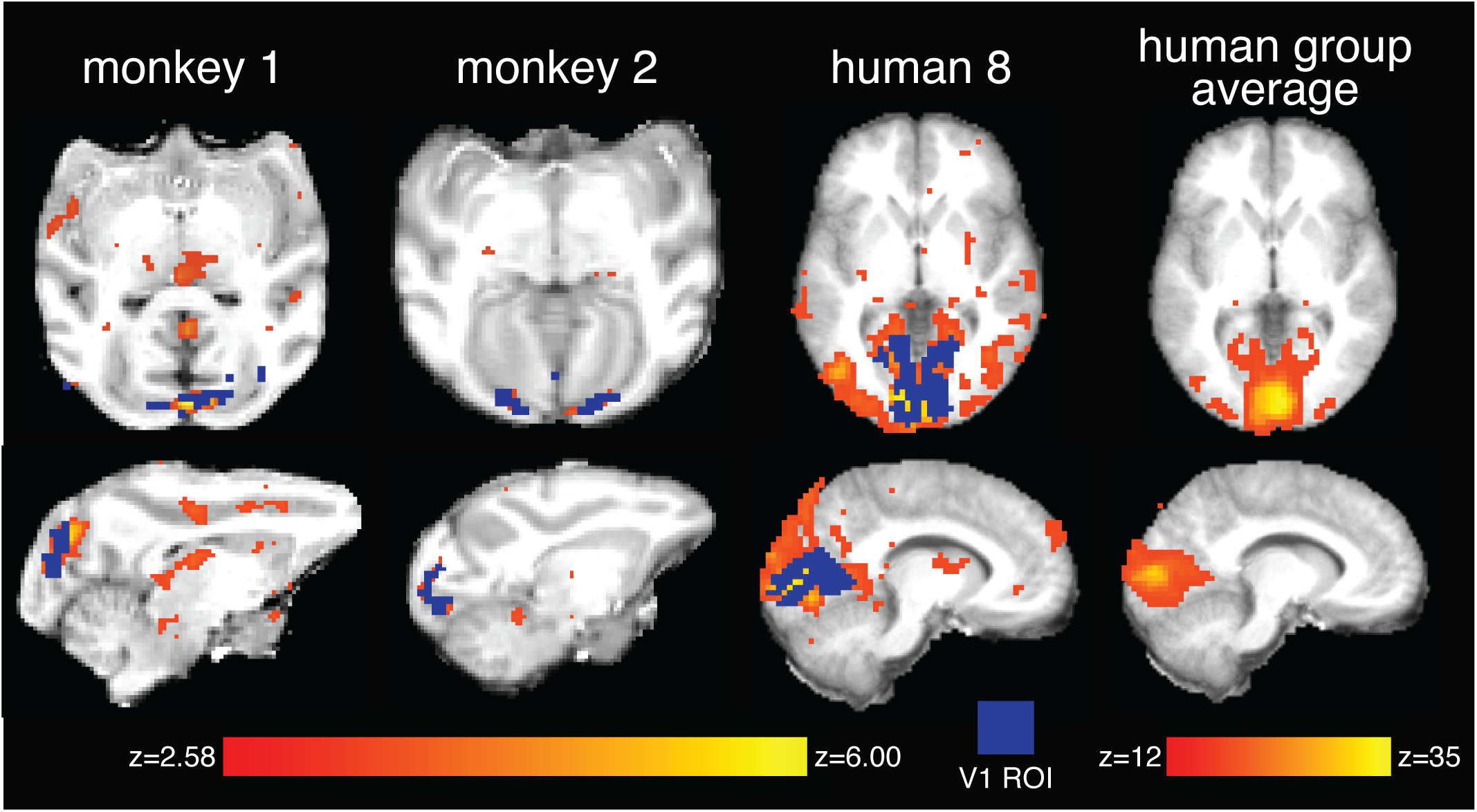
BOLD activation in visual cortex. Shown are transverse sections at the level of the calcarine sulci in both species. The three individual z-statistic maps came from an ANOVA of the main effect of time of the single-checkerboard response, and are thresholded at a p<.01 The group map is a random-effects average of the 12 humans, and is thresholded at p<10^−31^ to highlight the location of the activity peak. Voxels selected for the ROI from the 3-way conjunction are highlighted in blue in the individuals’ maps. See **Supplementary Figures** for whole brain coverage of the BOLD activations.

The next step was to isolate the BOLD response to each presentation of the second of the pair of checkerboards separated by one of the three ISIs. To do this, regressors were estimated for the linear trend, baseline, and for the 4 stimulus types (within-trial GLM: 1) single presentation plus all first checkerboard presentations, or second checkerboard presentations presented 2) 1.5 sec, 3) 3.0 sec, or 4) 4.5 sec after the first checkerboard). This regression procedure allowed for the separate estimation of the responses to the first and second checkerboards in the double-presentation trials, and has been previously validated and used in several recent human neuroimaging studies to separate activity related to different events within a trial (Shulman et al., 1999; Ollinger et al., 2001b; Ollinger et al., 2001a). Using this within-trial GLM, the baseline, linear trend, and single/first responses were removed from the original data set. For all subsequent analyses, these two new time-series of images were used, which contained the responses to 4 event-types: the single-presentation responses from the between-model data set, and the 3 second-presentation responses from the within-model data set. All of the GLMs and regression analyses were performed using in-house software.

Two analyses were performed on these extracted BOLD responses. The first analysis did not assume a BOLD response shape, allowing for any differences between the time-courses of the responses to be quantified. An ANOVA with trial as the repeated measure was used to determine in each subject if there was a significant effect of ISI x time between the single- and second-responses. A Greenhouse-Geisser correction was applied to correct the degrees of freedom for non-sphericity (SPSS, Chicago, Ill.). Two-tailed t-tests were performed comparing the value at each time-point of the single-presentation time-course versus each of the second-presentation time-courses. Comparisons were made between 0 and 10.5 sec after stimulus onset, which is the interval over which the single-presentation responses in the two monkeys and the human group average were above baseline.

In a second analysis, a hemodynamic response function was fit to the time-courses in order to quantify the differences in the amplitude of the responses at the different ISIs. First, a gamma function convolved with a .5 sec boxcar was fit (minimizing squared-error) to each subject’s single-response data, allowing τ to vary between .25 and 3.25 at .25 increments, and pure delay to vary between 1 and 6 seconds at 1 sec intervals (Boynton et al., 1996). Then this gamma function was used to fit the second response time-courses, allowing amplitude and delay to vary. Bootstrapping (resampling with replacement) was used to estimate the 95% confidence intervals of the mean amplitude and delay parameters of each of these fits; means were judged to be significantly different if the 95% confidence intervals did not overlap. Gamma function least-squares fits and bootstrapping were performed using MATLAB (Mathworks, Natick Mass.).

### Eye movement Analysis

The eye-tracking data from the two macaques were analyzed using custom software and the R statistical package (http://www.R-project.org). Putative eye movements were identified as changes in eye position of two degrees or more. The beginning, end, and peak velocity of these changes were then extracted, and main sequence relationships (duration versus peak velocity and duration versus average velocity) were used to separate true saccades from blink artifacts and measurement noise. Human eye movements were tracked with lower resolution, precluding a similar automated analysis. Instead, the data were visually inspected both on- and off-line to confirm that saccades were performed only rarely.

## Results

The present study was similar in design to that of Huettel and McCarthy (Huettel and McCarthy, 2000). In both species, subjects (2 monkeys, 12 humans) passively fixated a central point while either a single flickering checkerboard or a pair of flickering checkerboards (500 msec duration) with an inter-stimulus interval (ISI) of 1.5 sec, 3.0 sec, or 4.5 sec was presented on the screen (see **Figure 1**). In each subject, we defined a region in V1, and then the BOLD response to a single checkerboard from this region (**Figure 1B**) was compared with the response to the second of the pair of checkerboards at the 3 ISIs (**Figure 1C**). The results were then compared between the two species.

The two monkeys performed similarly, making 4.1 and 4.2 saccades of 2 deg or greater per minute, respectively. However, the distributions of the saccades were different: only 1.4% of the saccades in monkey 1 were initiated within 1.5s of the checkerboard presentation, whereas in monkey 2 this value was 31%. Human subjects made fewer than 1 medium or large saccade per minute.

In both monkeys, the presentation of a single checkerboard evoked activity in the calcarine sulci of both hemispheres— corresponding to the anatomical location of V1—as well as anatomical areas corresponding to extrastriate cortex and the MT/MST complex (see **Figure 2** and **Supplementary Figures 1** & **2**). In the human subjects, the activity was distributed in and around the calcarine sulci of both hemispheres, but in some cases also extended posteriorly toward the occipital pole, anteriorly toward the prostriate region, and to the lateral aspect of the occipital lobe (e.g. subject 8, **Figure 2**). When the activity was averaged over subjects, it was more focally centered along the calcarine sulcus, corresponding to the anatomical location of human V1 (human group average, **Figure 2** and **Supplementary Figure 3**). In humans, activity was also evoked in extrastriate areas and in the MT/MST complex. The human data had more statistical power as compared to the monkeys, because although we collected 3-4 times as many trials from the monkeys, the voxel size in the humans was much larger (19:1 volume ratio humans: monkeys). The resulting greater statistical power in the human data probably explains both why more voxels passed the statistical threshold for significance and why many voxels reached higher levels of significance in the humans as compared to the monkeys. For the statistical analysis carried out on the BOLD signal time-courses, we selected in each individual a region of interest (ROI) in area V1 that was defined both on anatomical (areal borders) and functional (presence of significant activation independently of any assumed HRF) criteria (the blue areas for individual subjects in **Figure 2**; see **Materials and Methods** for more detail on the ROI selection process).

Next, in both species, the BOLD responses to the second checkerboard at each of the three ISIs were separated from the responses to the pair of checkerboards. A general linear model (GLM) was used to estimate the response to the first checkerboard by modeling as one event type the single checkerboard trials and the first checkerboard of the double-checkerboard trials at all three ISIs. This combined estimate was then used to remove the first checkerboard response from each of the double-checkerboard responses, leaving behind the second checkerboard response at each of the three ISIs (within-trial model, see **Materials and Methods**). The validity of this model was confirmed in two different ways. First, the shape and magnitude of the response to the single checkerboard, when estimated in isolation (between-trial model), were not statistically distinguishable from estimates obtained by combining the response to the single checkerboard with the response to the first checkerboard in the pair (within-trial model). These results ensure that in the double presentation trials the estimation of the response to the second checkerboard was not contaminated by the response to the first checkerboard; if the single/first response had been different from the single response, this would have meant that some of the variance associated with the first response was mistakenly attributed to the second response. Second, subtracting the average response to the single checkerboards from the average response to the double checkerboard (shown schematically in **Figure 1**) yielded nearly identical estimates of the response to the second checkerboard as the regression procedure (with the exception of the 4.5 sec ISI time-course for monkey 1, which showed an earlier peak in the subtraction analysis). The latter method is analogous to `selective averaging’ as described in Dale and Buckner (Dale and Buckner, 1997), while the former method is derived from Shulman et al. (Shulman et al., 1999) and Ollinger et al. (Ollinger et al., 2001b; Ollinger et al., 2001a). Importantly, none of these models makes any assumption regarding the shape of the stimulus-evoked BOLD response.

**Figure 3:**
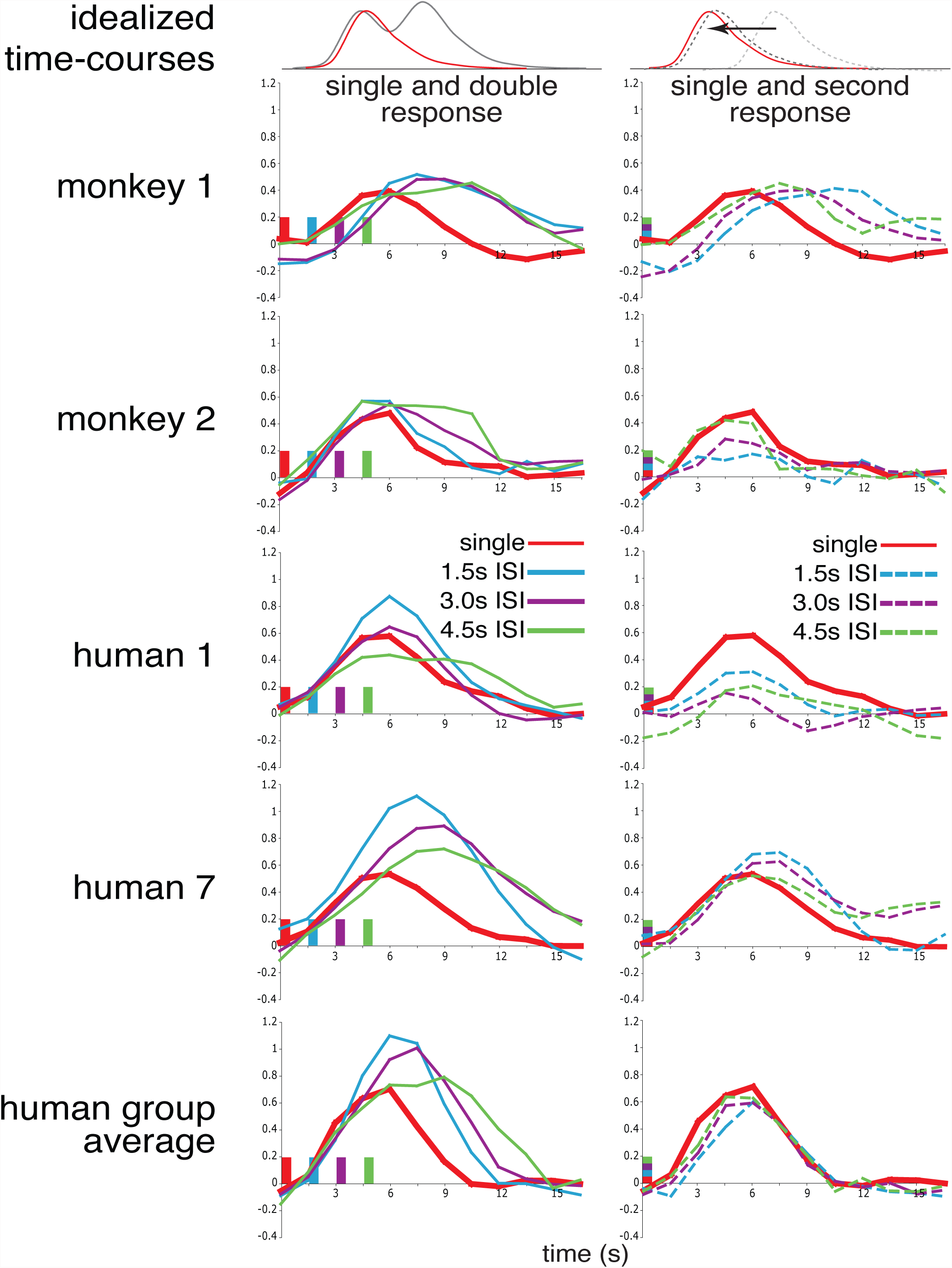
BOLD time-courses. *First column:* Average time-courses of activity evoked by a single checkerboard presentation compared to the activity evoked by the three trial types in which a pair of checkerboards separated by one of three ISIs (see legend) was presented. The red bar indicates the time of the checkerboard presentation in the single trials and the first checkerboard in the double trials; the other colors represent the times of the second checkerboard presentations in the double trials. *Second column:* Time-courses of the response to a single checkerboard compared to the response to the second of a pair of checkerboards after the first checkerboard response has been removed. The multi-colored bars indicate that in this figure the time-courses have been aligned on checkerboard presentation. For each row, the red “single” trace is the same in both columns. Time-courses in the top row are idealized BOLD responses illustrating the comparisons shown in the columns below.

**Figure 3** shows the time-course of the BOLD responses from both monkeys, two of the human subjects, and the average of all of the human subjects. The first column shows the average response to a single or double checkerboard at three different ISIs (1.5, 3.0, or 4.5 sec), as illustrated in **Figure 1B** and **1 A**, respectively. The second column shows the average response to a single checkerboard or to the second of a pair of checkerboards separated by one of the three ISIs, as illustrated in **Figure 1B** and **1C**, respectively. For ease of comparison, the individual time-courses have been smoothed with a 3 time-point binomial filter to limit high-frequency noise; all statistical tests were performed on unsmoothed data (see **Figure 4** for unsmoothed monkey time-courses). The individual human data (subjects 1 and 7) were chosen to illustrate human subjects with BOLD responses that are similar to those of monkey 2 and monkey 1, respectively. Human 1, like monkey 2, demonstrates suppression of the second BOLD response even at the longest ISI of 4.5 sec. Human 7, like monkey 1, demonstrates no suppression of the second BOLD response at any ISI, and also a delay in the time to peak.

We first analyzed whether the length of the ISI influenced the shape of the BOLD response. In both monkeys, there was a significant effect of ISI length on the response shape (monkey 1: ANOVA(ISI) d.f.=3 F=3.633 p<.05; monkey 2: ANOVA(ISI) d.f.=3 F=3.370 p<.05). In general, at the shortest ISI (1.5 sec) the shape of the second checkerboard response differed significantly from the single checkerboard response, and this difference disappeared at the longest ISI (4.5 sec). This effect can be seen in **Figure 3** as well as in the results of t-tests performed at each time-point (**Table 1**). These t-tests indicate that not only was there a significant effect of ISI, but that this effect was only present at certain time-points. In both animals, at 1.5 sec ISI the second response differed significantly (p<.05) from the single response at several time points corresponding to the upslope and peak (monkey 1: time points between 3-4.5 sec; monkey 2: between 4.5-6 sec). In contrast the 4.5 sec ISI response was not significantly different from the single response in either monkey near the peak. This interaction of ISI and time was confirmed in both monkeys by an ANOVA (monkey 1: ANOVA(ISI,time) d.f.=10.240 F=4.464 p<.0001; monkey 2: (ANOVA(ISI,time) d.f.=10.458 F=2.683 p<.01).

**Table 1:**
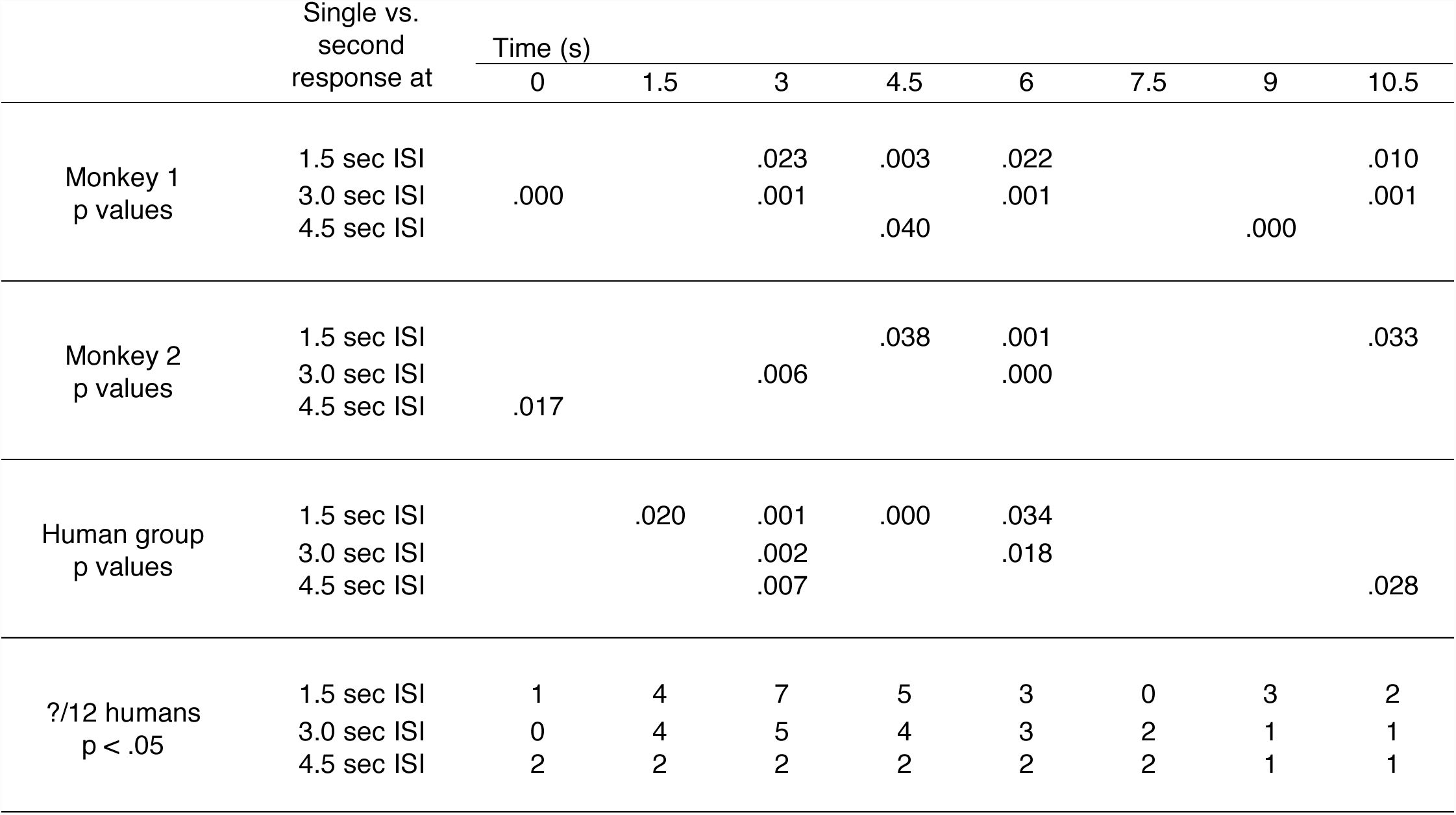
Comparison of single- and second-presentation response time-courses. Shown are the p-values from the t-tests (not multiple comparisons corrected) comparing the single-presentation BOLD response with that of the second-presentation time-courses at the 3 ISIs for each time-point between 0 and 10.5 sec. *Sections 1 and 2:* Time-points where the comparison was significant (p<.05) within each individual monkey. *Section 3:* Time-points from the human data averaged across the group where the comparison was significant (p<.05). *Section 4:* The number of human subjects at each time-point (out of 12) where the comparison was significant (p<.05). Time-points at which the second response differs from the single response

|  | Single vs.<br>second<br>response at | Time (s) |  |  |  |  |  |  |  |
| --- | --- | --- | --- | --- | --- | --- | --- | --- | --- |
|  |  | 0 | 1.5 | 3 | 4.5 | 6 | 7.5 | 9 | 10.5 |
| Monkey 1<br>p values | 1.5 sec ISI |  |  | .023 | .003 | .022 |  |  | .010 |
|  | 3.0 sec ISI | .000 |  | .001 |  | .001 |  |  | .001 |
|  | 4.5 sec ISI |  |  |  | .040 |  |  | .000 |  |
| Monkey 2<br>p values | 1.5 sec ISI |  |  |  | .038 | .001 |  |  | .033 |
|  | 3.0 sec ISI |  |  | .006 |  | .000 |  |  |  |
|  | 4.5 sec ISI | .017 |  |  |  |  |  |  |  |
| Human group<br>p values | 1.5 sec ISI |  | .020 | .001 | .000 | .034 |  |  |  |
|  | 3.0 sec ISI |  |  | .002 |  | .018 |  |  |  |
|  | 4.5 sec ISI |  |  | .007 |  |  |  |  | .028 |
| ?/12 humans<br>p < .05 | 1.5 sec ISI | 1 | 4 | 7 | 5 | 3 | 0 | 3 | 2 |
|  | 3.0 sec ISI | 0 | 4 | 5 | 4 | 3 | 2 | 1 | 1 |
|  | 4.5 sec ISI | 2 | 2 | 2 | 2 | 2 | 2 | 1 | 1 |

In humans, there was also a significant effect of ISI and ISI x time on the time-course of the BOLD response in 7 of the 12 individuals (for each individual, ANOVA(ISI, time) p<.05 or ANOVA(ISI) p<.05), as well as in the group average (ANOVA(ISI) d.f.=2.31 F=3.814 p<.05; ANOVA(ISI, time) d.f.=6.69 F=2.085 p=.0584). The time-point paired t-tests across conditions (third section, **Table 1**) shows that at the 1.5 sec ISI, the group average response to the second checkerboard was significantly different from the single checkerboard response at all time-points between 1.5 and 6 sec after the onset of the stimulus. In contrast, at the 4.5 sec ISI, the time-course differed only at 3.0 sec.

However, underlying these group average results in the humans was a high degree of individual variability. Across the 12 subjects, there were substantial departures from the group average (e.g., **Figure 3**, rows 4 and 5). This variability is summarized in the fourth section of **Table 1**, which shows the number of individual human subjects (out of 12) demonstrating significant departures from linearity at each time-point for each ISI condition. At each time-point in which there was a significant difference between the single and second response in the group average, at least 5 subjects failed to show a significant difference. Conversely, at time-points in which there were no significant differences in the group average, at many as 3 subjects still demonstrated significant differences between the single and second responses.

Despite the high degree of variability between individuals, for each increasing step in ISI length, fewer subjects demonstrated a significant difference between the second and single response during the upslope and peak of the response (1.5 to 4.5 seconds after stimulus onset). For the 1.5 sec ISI condition, up to 7 out of 12 subjects demonstrated a significant difference at each time-point between 1.5 and 6 sec after stimulus onset. For the 3.0 sec ISI condition, only 5 of 12 subjects showed significant difference at any single time-point. Finally, for the 4.5 sec ISI condition, at the only time-point in which the group average time-course exhibited a significant difference, only 3 of the 12 subjects actually demonstrated a difference. This pattern of results from the human group indicates that the variability between the two monkeys in the response time-courses is not inconsistent with the variability seen in the humans.

Another way to assess response variability is to compare the human and monkey BOLD time-courses at each ISI. This can be seen in **Figure 4**, which compares the two monkeys response time-courses to the 1 and 2 standard deviation ranges of the human group. In general, the monkey data fall within 2 standard deviations of the human mean. The exception is the 1.5 sec ISI time-course, where both monkeys’ time-courses are more delayed and sustained than the humans.

**Figure 4:**
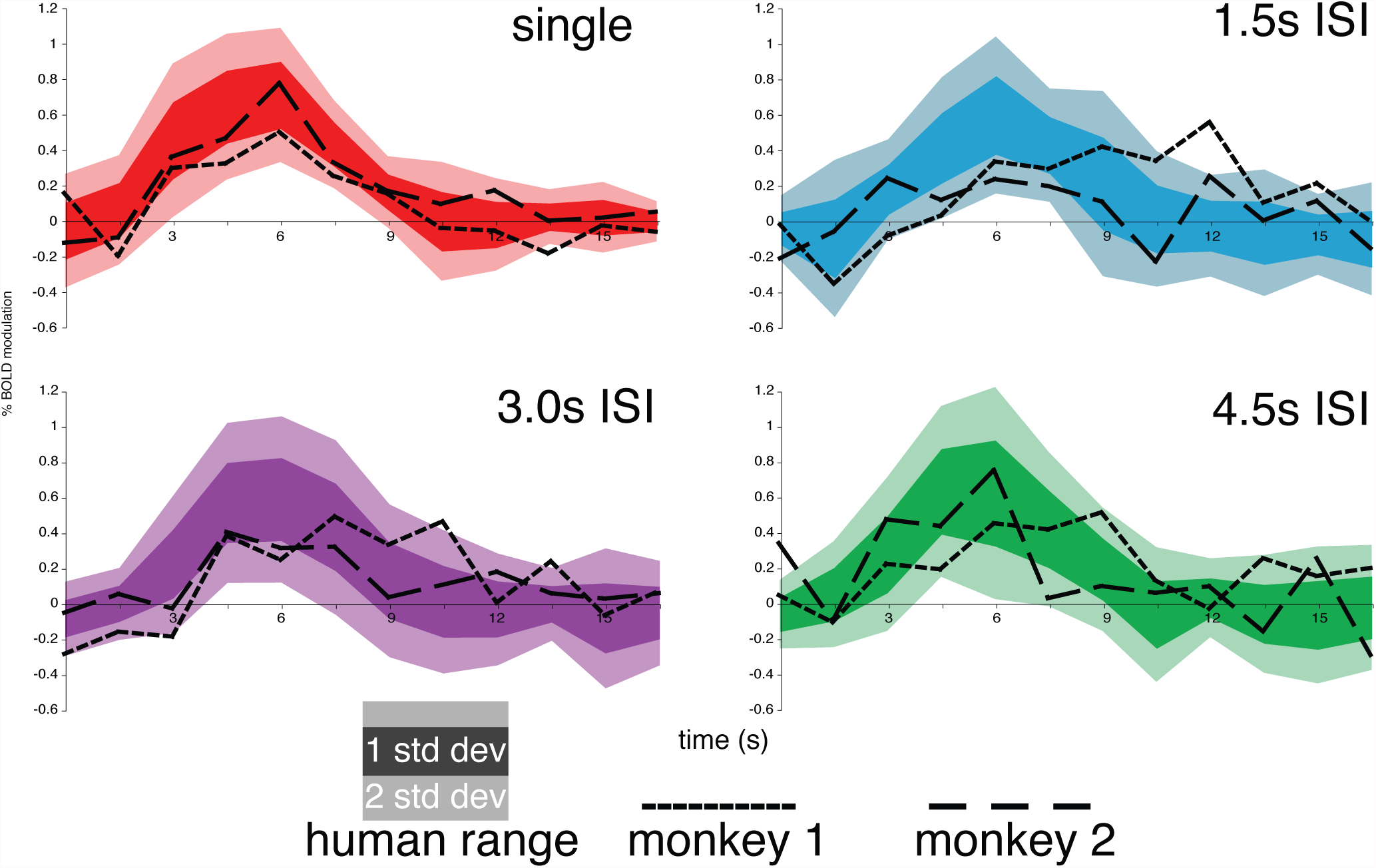
Monkey individuals vs. human group variability. In each panel, time-courses of the response to a single- or second-presentation of a checkerboard of the two monkeys (unsmoothed) are plotted over the variation in the human group data. The dark band of color represents 1 standard deviation of the human group mean, and the light band of color represents 2 standard deviations.

In a second analysis, we fit a gamma function to the time-courses to measure the magnitudes of the response to the single and second checkerboards. We first fit a gamma function to the mean time-course of the BOLD response to the single checkerboard in order to compare the magnitude and shape of the single response between the two species. The magnitudes of each of the two monkeys’ responses (.46% and .57% BOLD modulation) fell within 2 standard deviations of the human mean (.72% +/− .16%). In addition, the other parameters of the fitted gamma functions were similar across the two species (monkey 1: delay=1.82 sec, tau=1.25; monkey 2: delay=1.26 sec, tau=1.75; human group mean +/− 1 SD: delay=1.46 +/− .74 sec, tau=1.46 +/− .33).

We then fit the gamma function derived from the single-checkerboard response to the second checkerboard BOLD responses at each of the three ISIs. To compare linearity as a function of ISI among individuals and between species, we normalized the results for each individual by expressing the response magnitude to the second checkerboard as a fraction of the response magnitude to the single checkerboard (see **Figure 5**). In both monkeys, the magnitude of the BOLD signal to the second checkerboard increased with the length of the ISI, although this effect was significant only in monkey 2 (1.5 sec vs. 4.5 sec ISI, monkey 2, bootstrap, p<.01). The human group data also demonstrated stronger responses to the second checkerboard as function of ISI length (one-tailed paired t-test, p<.05). In both monkeys, the magnitude of the BOLD signal at the 4.5 sec ISI recovered to between 80% and 120% of the single checkerboard BOLD signal (monkey 1: 116%; monkey 2: 86%). These values were within one standard deviation of the human group mean (95 +/− 31%, n=12). At the shorter ISIs, the two monkey’s ratios remained within 1-2 standard deviations of the human mean, though monkey 2 was trending away from the human mean at the 1.5 sec ISI.

Although the smaller voxels in the monkey data resulted in less statistical power per voxel, the results from each monkey are highly reliable. We tested within-subject reliability in two ways. First, we split the total data set from each monkey into half, according to whether it was collected in the first (early) or second (late) half of the scanning sessions. We then repeated the gamma fitting procedure on each half. The results of this are shown in **Figure 6** in the orange and green lines. In each monkey, the results from each half of the data were consistent with the mean result from the entire data set (shown in black), and the results from the two monkeys did not overlap except at the longest ISI. Second, we assessed the reliability of the mean with bootstrapping. The error bars in Figure 6 show the 95% confidence intervals derived from this test. The BOLD response amplitude at all three ISIs in monkey 1 was statistically indistinguishable from the single response. In monkey 2, the amplitude of the second response was significantly suppressed as compared to the single response at the 1.5 sec ISI, but not at the 3.0 and 4.5 sec ISIs.

**Figure 5:**
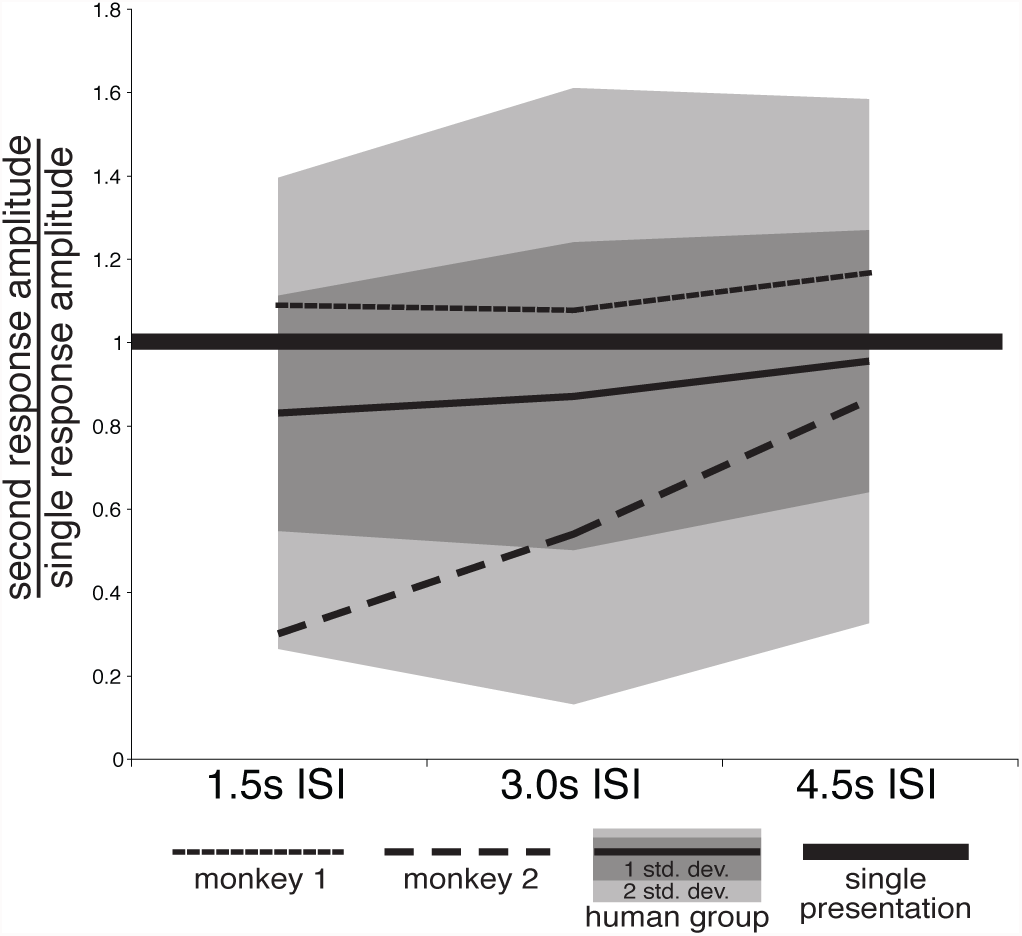
Second vs. single response ratios at the three ISIs. For both monkeys and the human group, the graph shows the ratio of the amplitude of the second-presentation BOLD responses after a 1.5, 3.0, or 4.5 sec ISI divided by the amplitude of the single BOLD response. A ratio of 1.0 means the amplitude of the second response equals that of the single response. The shading outlines the range of 1 (dark) and 2 (light) standard deviations from the human group mean.

**Figure 6:**
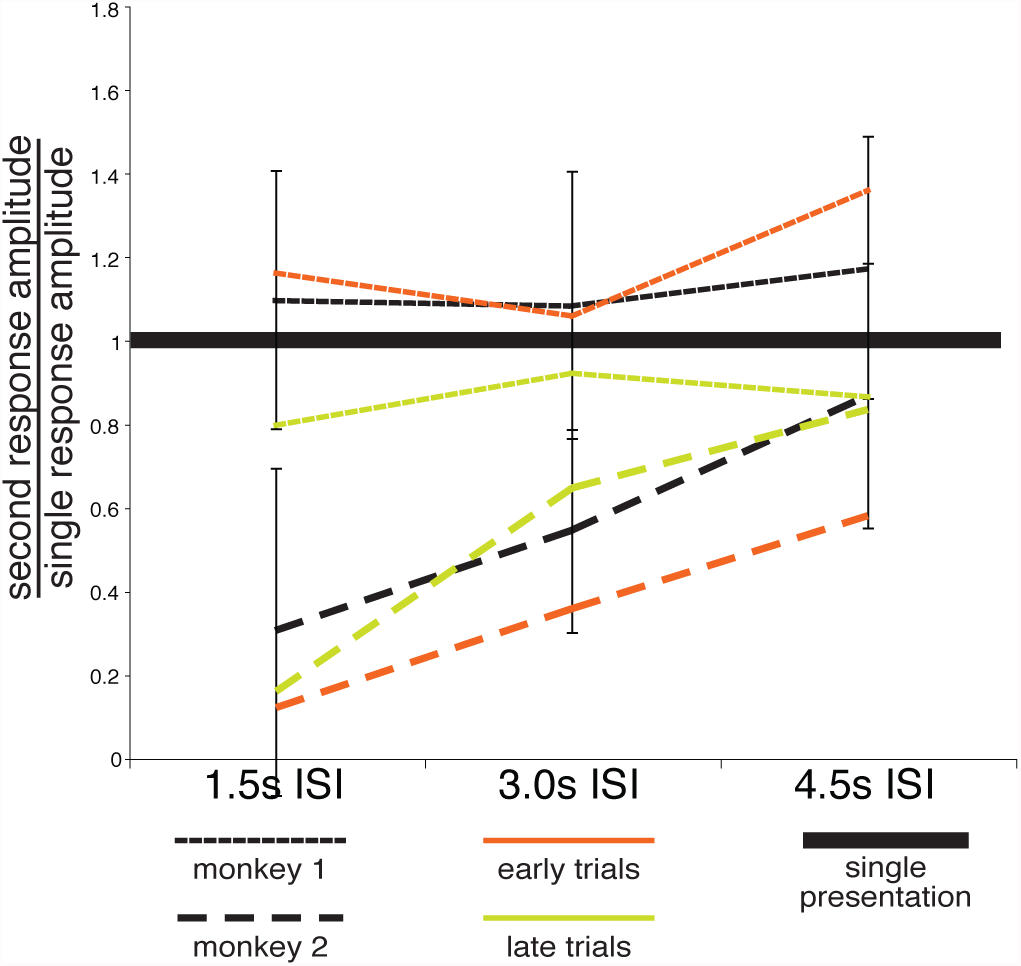
Reproducibility of the monkey data. The orange and green lines represent the amplitude ratios resulting from repeating the gamma-function fitting procedure on each half of the data. The error bars represent the standard error of the mean derived from a bootstrap test.

## Discussion

We compared several parameters of the BOLD response (shape, magnitude, and summation of the overlapping responses) to closely spaced events in the visual cortex of human and macaque subjects during the presentation of flickering checkerboards. We found that the stimulus-evoked BOLD responses in primary visual cortex were largely similar across the two species.

Other studies have described the shape and magnitude of the BOLD response in macaques to periods of continuous stimulation in fMRI blocked designs or to single motor or cognitive events. Logothetis *et al.* have found that the BOLD responses in anesthetized macaques have the same general shape as humans, and that, as in humans, the transformation of neural activity to a BOLD response is linear (Logothetis et al., 2001). More recently Leite and Mandeville determined that BOLD response detection power is essentially unchanged at ISIs varying from 4 to 20 seconds, suggesting that in that range the BOLD response summation is probably linear (Leite and Mandeville, 2006). However, they used long stimulation durations (4 seconds) and modeled the hemodynamic response as two exponential functions; these two decisions make comparisons to the extant human literature difficult, since the human studies have generally used very short stimulus durations (.5-1 seconds) and gamma functions as the model hemodynamic response (Leite et al., 2002; Leite and Mandeville, 2006).

Several studies have also directly compared human and macaque fMRI data. Two studies presented stimuli spaced by the same range of ISIs in both humans and monkeys in a rapid event-related design, with the shortest ISI being 2 sec; these studies implicitly assumed that BOLD responses added similarly in the two species (Nakahara et al., 2002; Koyama et al., 2004). Koyama *et al.* noted that in putatively homologous oculomotor areas, the macaque BOLD responses during visually-guided saccades in both blocked- and event-related experiments were larger in magnitude than that of humans (Koyama et al., 2004). However, it is unclear whether these differences arise from real interspecies physiological differences, from behavioral differences, or from experimental differences between the two species. Denys *et al.* compared the magnitudes of responses to stimuli presented in a block design and found a large difference between the two species, but attributed this result to differences in volitional control of stimulus processing. Further complicating the human-monkey comparison of the response magnitude was the fact that they were comparing monkey hemodynamic responses measured with a contrast agent (MION) with human BOLD responses—MION and BOLD responses have been shown to have different shapes and detection power (Vanduffel et al., 2001; Denys et al., 2004b). Overall, across experiments, the presence of either potentially significant methodological or behavioral differences during complex tasks have so far prevented any firm conclusion on the comparability of stimulus-evoked BOLD responses in monkey and humans.

Notwithstanding that behavioral strategy or levels of attention may be nearly impossible to control, our strategy was to compare BOLD responses in human and macaque primary visual cortex to briefly presented and powerful visual stimuli. This approach should minimize potential behavioral or functional-anatomical differences as it is generally agreed that the organization and function of primary visual cortex or area V1 are largely similar in the two species (Van Essen et al., 2001; Tootell et al., 2003; Orban et al., 2004); that the influence of attention on V1 neuronal activity is relatively small as compared to higher order visual areas (Motter, 1993; Luck et al., 1997; Gandhi et al., 1999; Somers et al., 1999); and that neural responses to the passive presentation of high contrast stimuli is weakly or not at all modulated by attention (Reynolds and Desimone, 2003).

There are two main findings from the single checkerboard presentations. First, the magnitude of the visually evoked BOLD response to a 500ms high contrast flickering checkerboard was slightly lower in macaque than human visual cortex (monkey 1, .46%; monkey 2, .57%; human mean, .72%), but well within the individual human variability (2 std. dev. = .31%). Second, the shape of the BOLD response, as quantified by the parameters of the fitted gamma functions, was similar across the two species (monkey 1 delay=1.82 sec, monkey 2 delay=1.26 sec, human group mean (std. dev.) delay=1.46 sec (.74); monkey 1 tau=1.25, monkey 2 tau=1.75, human group mean (std. dev.) tau=1.46 (.33)). The shape parameters in both species agree with those of previous human studies (Boynton et al., 1996; Aguirre et al., 1998). These findings suggest that at least in primary visual cortex, potential stimulus-evoked BOLD response differences between humans and monkeys cannot be explained by differences in neurovascular coupling or the physiology of the BOLD signal, hence validating the use of BOLD fMRI for comparing visual response magnitudes in the two species. If we generalize these results to higher-order areas, we have to infer that previously reported differences in response magnitude during more complex tasks and higher order areas (Denys et al., 2004b; Koyama et al., 2004) are not due to differences in BOLD physiology, but rather reflect behavioral or neural differences.

When we presented two flickering checkerboards separated by a variable ISI (1.5, 3.0, and 4.5 seconds), we found that BOLD responses in visual cortex added nearly linearly and to a similar extent in human and macaque subjects. In both species at a 4.5 sec ISI, the magnitude of the response to the second stimulus was nearly the same as that of the single stimulus (second response/single response: monkeys 86-116%, humans 95% +/− 31%). This result indicates near linearity of BOLD response summation at this ISI. As the ISI decreased in length, individuals of both species showed an increased likelihood of having a decreased second checkerboard response amplitude.

These results are in keeping with previous work in humans (Dale and Buckner, 1997; Huettel and McCarthy, 2000; Miezin et al., 2000; Boynton and Finney, 2003). While Huettel and McCarthy demonstrated that even at an ISI of 6 sec, BOLD responses in humans do not add linearly, other studies have found that in V1 BOLD responses can add nearly linearly with ISIs as short as 2 seconds (Huettel and McCarthy, 2000). Miezin *et al.* found that while using a jittered ISI of 5 +/−2.5 sec did cause a small but significant reduction in response amplitude, the departures from linearity did not cause a significant decrease in the ability to detect a hemodynamic response (Miezin et al., 2000). Overall our findings indicate that, for cross-species studies using rapid event-related designs, events should be spaced at least 4.5 seconds apart to obtain comparable estimates and linear summation in the two species.

Our results also indicate substantial inter-subject variability in both human and monkey subjects in terms of shape and magnitude of the BOLD response (**Figure 4**, **Table 1**) and the linearity of BOLD response summation (**Figures 5** and **6**). This fact has not always been clearly highlighted in the published human studies where results are typically presented as across-subject group averages. In our human group, the across-subject mean signal time course for single or double stimuli is entirely consistent with what reported in prior work. However, individually, there were extreme variations consistent with research showing that between-subject variability is 16 times larger than within-subject variability (Aguirre et al., 1998). Some subjects showed slightly higher (rather than lower) signal magnitude to the second checkerboard, while others showed a large decrease in the second checkerboard response amplitude, even at the 4.5 sec ISI (see subjects 1 and 7 in **Figure 3**). While we recognize that with only two subjects tested, we have clearly not fully characterized inter-subject variability in macaque, we nonetheless observe that the variability in these two monkeys was not outside of the human range.

Establishing the minimal ISI at which the BOLD responses add linearly in both species is important for two reasons. ISIs that are too short may substantially reduce the magnitude of responses to closely spaced stimuli, which in turn will reduce the sensitivity for detecting responses. This possibility of reduced sensitivity is particularly worrisome in the monkey, since it will compound the already relatively low power of the monkey fMRI technique, which results mainly from the smaller voxels (1.5 mm vs. 3-4 mm in human fMRI) that are typically used for acquisition. Moreover, differences in BOLD non-linearities between species, which may be especially severe at the shorter ISIs, may confound inter-species comparisons of response magnitude. This will be true in experimental designs that do not allow for proper counter-balancing of event types, such as delayed-saccade or match-to-sample paradigms in which two stimuli must be presented in a fixed sequence (Aguirre et al., 1998), or in designs in which the response magnitude to the second of a pair of stimuli is the main dependent variable (e.g. adaptation paradigms, (Grill-Spector and Malach, 2001).

Despite our attempts to make the monkey and human experiments as similar as possible, several aspects of data collection still differed, and may have caused differences in the resulting BOLD responses. Artifactual differences may be introduced by mouth movements related to consuming a reward, differences in skull geometry, voxel size, and the presence of head-restraint devices in the monkeys. In addition to these experimental differences, there may have been inter-species differences in the vasculature, in the neural activity evoked by a stimulus, and in the relationship between the neural activity and the BOLD response (Cannestra et al., 1998; Buxton et al., 2004; Fox et al., 2005). Given all of these factors, it is remarkable that the properties of the BOLD response in visual cortex do not vary more between humans and macaques.

In conclusion, the response characteristics of visually evoked BOLD signals appear to be similar in monkey and human visual cortex, and accurate response magnitudes can be estimated by spacing these stimuli at least 4.5 seconds apart. This information can now be used to guide the design of future cross-species event-related fMRI protocols, and also increase the confidence in the results of such studies.

## Acknowledgements

We would like to thank Roger Tootell and Wim Vanduffel for their advice on developing the experimental setup, Larry Wald for the MR coils, and Avi Snyder, Mark McAvoy, Anthony I. Jack, and Giovanni D’Avossa for their advice on the analysis and comments on the manuscript. This work was generously supported by the McDonnell Center for Higher Brain Function, the Washington University Conte Center, and the NIH (NINDS, NEI, NIMH). Address correspondence to Gaurav H. Patel, Dept. of Radiology, Washington University School of Medicine, 4525 Scott Ave., Room 2123, St. Louis, MO 63110.

---

**Supplementary Figure 1.**
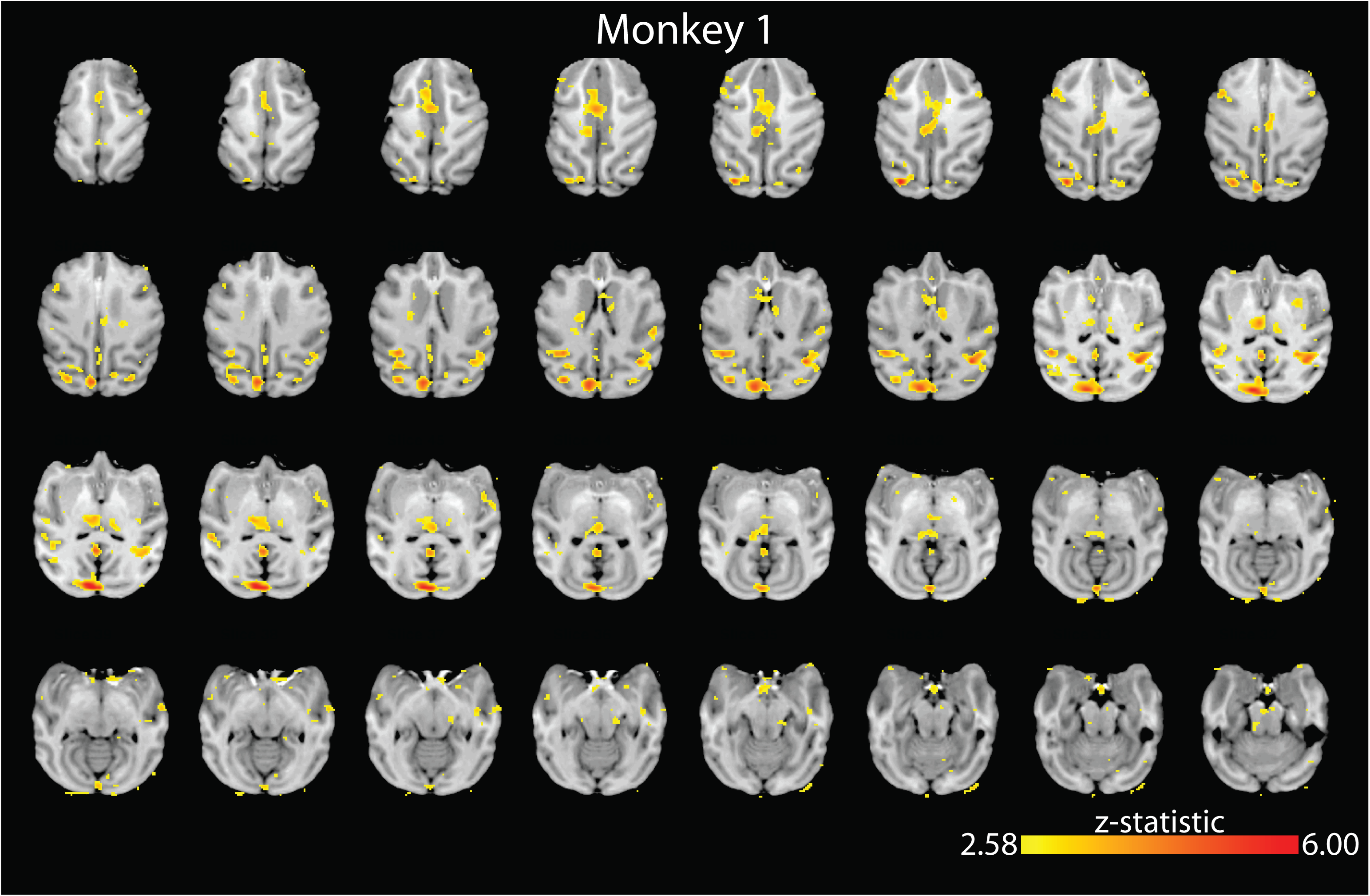

**Supplementary Figure 2.**
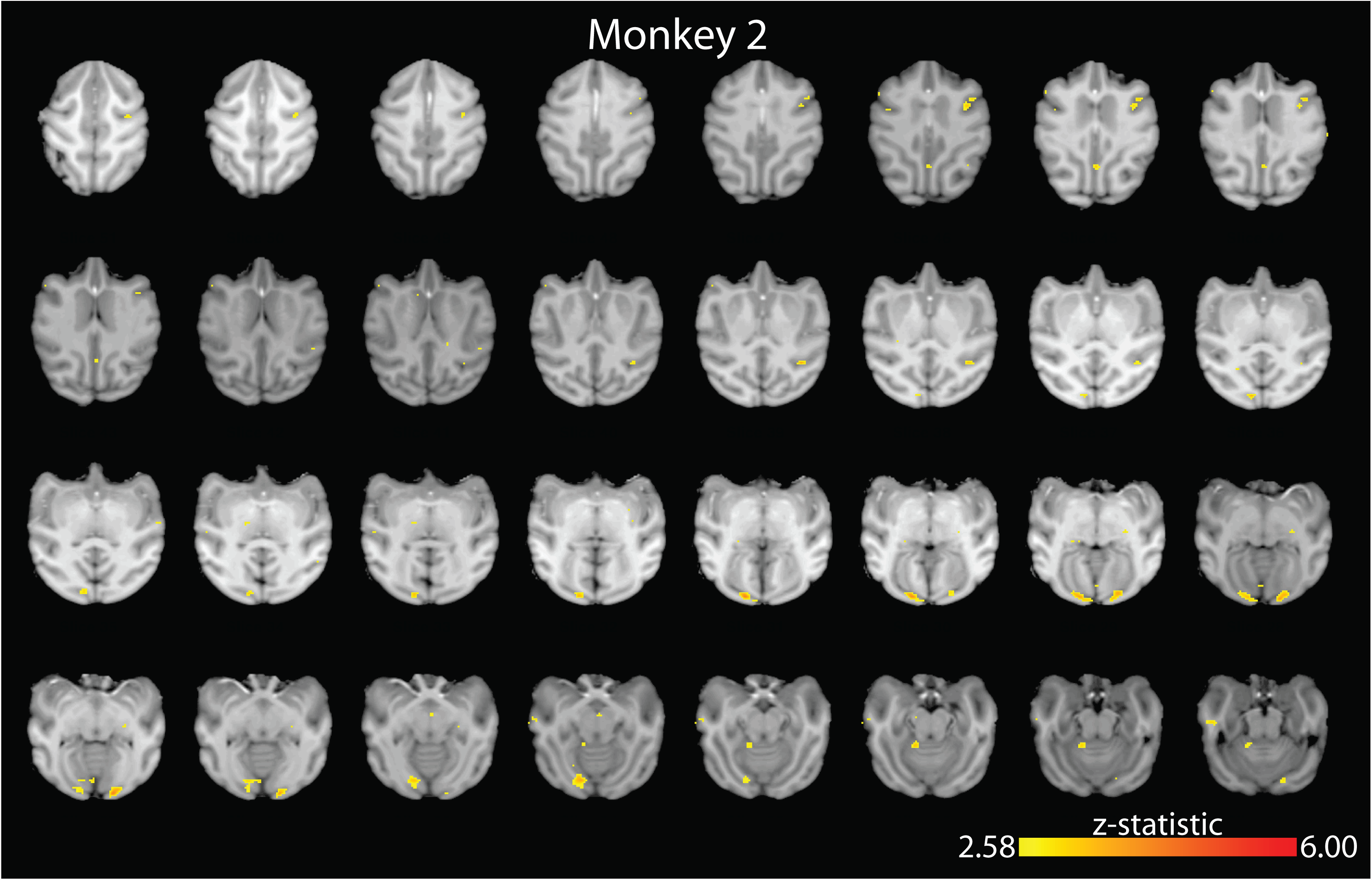

**Supplementary Figure 3.**
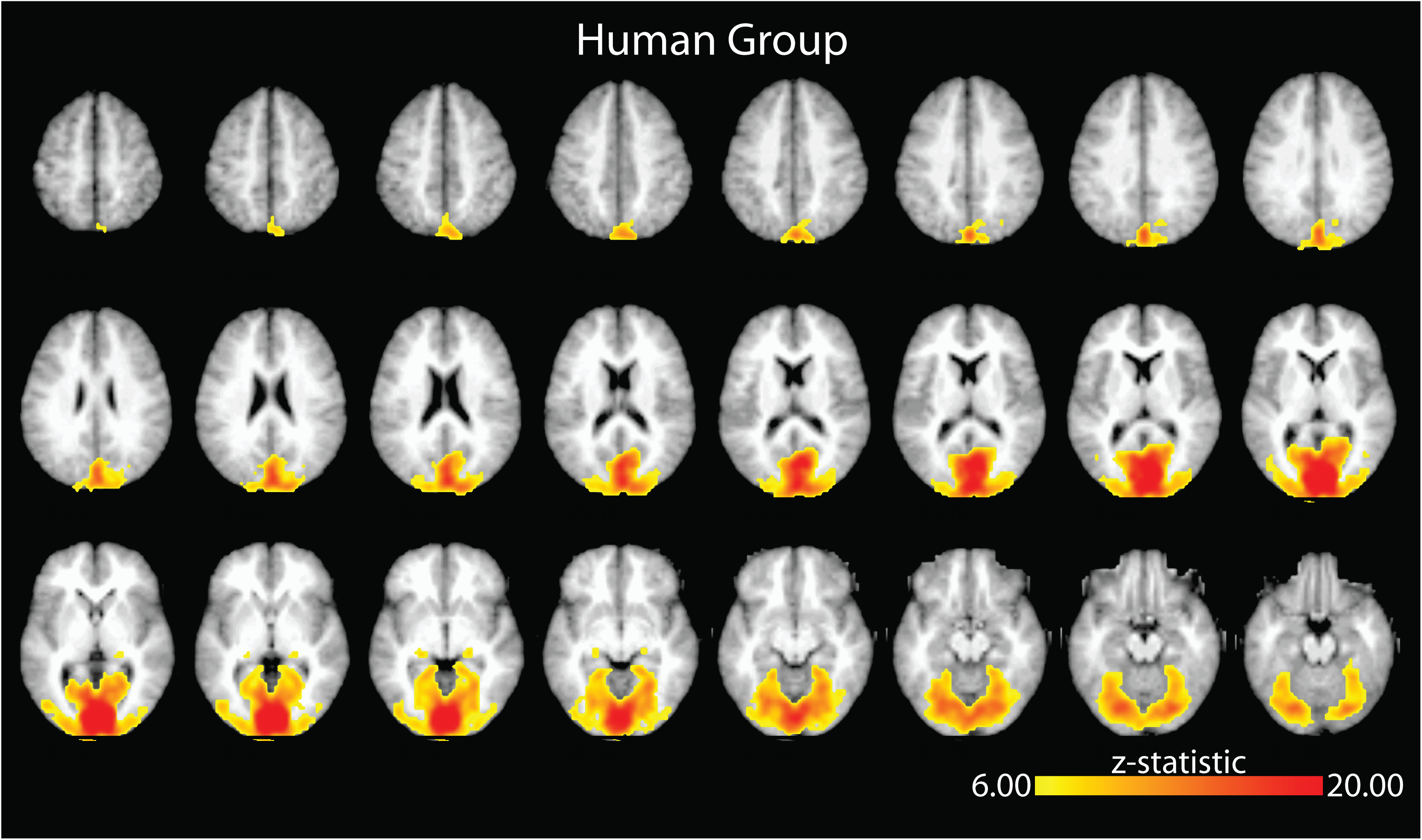

